# The BK channel gating ring is strongly coupled to the voltage sensor

**DOI:** 10.1101/381319

**Authors:** Pablo Miranda, Miguel Holmgren, Teresa Giraldez

## Abstract

The open probability of large conductance voltage- and calcium-dependent potassium (BK) channels is regulated allosterically by changes in the transmembrane voltage and intracellular concentration of divalent ions (Ca^2+^ and Mg^2+^). The divalent cation sensors reside within the gating ring formed by eight Regulator of Conductance of Potassium (RCK) domains, two from each of the four channel subunits. Overall, the gating ring contains 12 sites that can bind Ca^2+^ with different affinities. Using patch-clamp fluorometry, we have shown robust changes in FRET signals within the gating ring in response to divalent ions and voltage, which do not directly track open probability. Only the conformational changes triggered through the RCK1 binding site are voltage-dependent in presence of Ca^2+^. Because the gating ring is outside the electric field, it must gain voltage sensitivity from coupling to the voltage-dependent channel opening, the voltage sensor or both. Here we demonstrate that alterations of voltage sensor dynamics known to shift gating currents produce a cognate shift in the gating ring voltage dependence, whereas changing BK channels’ relative probability of opening had little effect. These results strongly suggest that the conformational changes of the RCK1 domain of the gating ring are tightly coupled to the voltage sensor function, and this interaction is central to the allosteric modulation of BK channels.

## INTRODUCTION

The open probability of large conductance voltage-and Ca^2+^-activated K^+^ (BK or slo1) channels is regulated allosterically by voltage and intracellular concentration of divalent ions (1–6). This feature makes BK channels important regulators of physiological processes such as neurotransmission and muscular function, where they couple membrane voltage and the intracellular concentration of Ca^2+^ (7–10). The BK channel is formed in the membrane as tetramers of α subunits, encoded by the KCNMA1 gene (11, 12). Each α subunit contains seven transmembrane domains (S0 to S6), a small extracellular N-terminal domain and a large intracellular C-terminal domain (13–15) (**Fig. 2a**). Similar to other voltage-gated channels, the voltage across the membrane is sensed by the voltage sensor domain (VSD), containing charged amino acids within transmembrane segments S2, S3 and S4 (15–20). The sensor for divalent cations is at the C-terminal region and is formed by two Regulator of Conductance for K^+^ domains (RCK1 and RCK2) per α subunit (2, 4, 15, 21–23). In the tetramer, four RCK1-RCK2 tandems pack against each other in a large structure known as the gating ring (15, 21, 24). Two high-affinity Ca^2+^ binding sites are located in the RCK2 (also known as “Ca^2+^ bowl”) and RCK1 domains, respectively. Additionally, a site with low affinity for Mg^2+^ and Ca^2+^ is located at the interface between the VSD and the RCK1 domain (4, 15, 25, 26) (**Fig. 2a**). The high-affinity binding sites show structural dissimilarity (15, 27) and different affinity for divalent ions (2). Apart from Ca^2+^, it has been described that Cd^2+^ selectively binds to the RCK1 site, whereas Ba^2+^ and Mg^2+^ show higher affinity for the RCK2 site (2, 4, 25, 26, 28–30). Thus, intracellular concentrations of Ca^2+^, Cd^2+^, Ba^2+^ or Mg^2+^ can shift the voltage dependence of BK activation towards more negative potentials. Using patch clamp fluorometry (PCF), we have shown that these cations trigger independent conformational changes of RCK1 and/or RCK2 within the gating ring, measured as large changes in the efficiency of Fluorescence Resonance Energy Transfer (FRET) between fluorophores introduced into specific sites in the BK tetramer. These rearrangements depend on the specific interaction of the divalent ions with their high-affinity binding sites, showing different dependences on cation concentration and membrane voltage (30, 31). To date, the proposed transduction mechanism by which divalent ion binding increases channel open probability was a conformational change of the gating ring that leads to a physical pulling of the channel gate, where the linker between the S6 transmembrane domain and the RCK1 region acts like a passive spring (32). Such a mechanism would be analogous to channel activation by ligand binding in glutamate receptor or cyclic nucleotide-gated ion channels, also tetramers (33, 34). Our previous results do not support this as the sole mechanism underlying coupling of divalent ion binding to channel opening, since the gating ring conformational changes that we have recorded: 1) are not strictly coupled to the opening of the channel’s gate, and 2) show different voltage dependence for each divalent ion. In addition, the recent cryo-EM structure of the full slo1 channel of *Aplysia californica* (15, 22) shows that the RCK1 domain of the gating ring is in contact with the VSD, predicting that changes in the voltage sensor position could be reflected in the voltage dependent gating ring reorganizations.

Understanding the nature of the voltage dependence associated to individual rearrangements produced by binding of divalent ions to the gating ring is essential to untangle the mechanism underlying the role of such rearrangements in BK channel gating. To this end, we now performed PCF measurements with BK channels including a range of VSD mutations or coexpressed with different regulatory subunits. Here we provide evidence for a functional interaction between the gating ring and the voltage sensor in full-length, functional BK channels at the plasma membrane, in agreement with the structural data from *Aplysia* BK. Moreover, these data support a pathway that couples of divalent ion binding to channel opening through the voltage sensor.

## RESULTS

### Voltage dependence of gating ring rearrangements is associated to activation of the RCK1 binding site

BK α subunits labeled with fluorescent proteins CFP and YFP in the linker between the RCK1 and RCK2 domains (position 667) retain the functional properties of wild-type BK channels (30, 31). This allowed us to use PCF to detect conformational rearrangements of the gating ring measured as changes in FRET efficiency (*E*) between the fluorophores (30, 31). Binding of Ca^2+^ ions to both high-affinity binding sites (RCK1 and Ca^2+^ bowl) produces an activation of BK channels, coincident with an increase in *E* from basal levels reaching saturating values at high Ca^2+^ concentrations ((31) and **Fig. 1a**). In addition, we observed that the *E* signal has the remarkable property that in intermediate Ca^2+^ concentrations it shows voltage dependence besides its Ca^2+^ dependence ((31) and **Fig. 1a**). Independent activation of high-affinity binding sites by other divalent ions (Ba^2+^, Cd^2+^, or Mg^2+^ (30)) led us to postulate that Ca^2+^ activation has a site-dependent relation to voltage. To further evaluate the effect of individual high-affinity Ca^2+^ binding sites on the voltage-dependent component of the gating ring conformational changes we first selectively mutated the binding sites. Mutations D362A and D367A (2, 4) were introduced in the BK667CY construct (BK667CY^D362A/D367A^) to remove the high-affinity binding site located in the RCK1 domain. **Fig. 1b** shows the relative conductance and *E* values for the BK667CY^D362A/D367A^ construct at different membrane voltages for various Ca^2+^ concentrations.

**Fig. 1.**
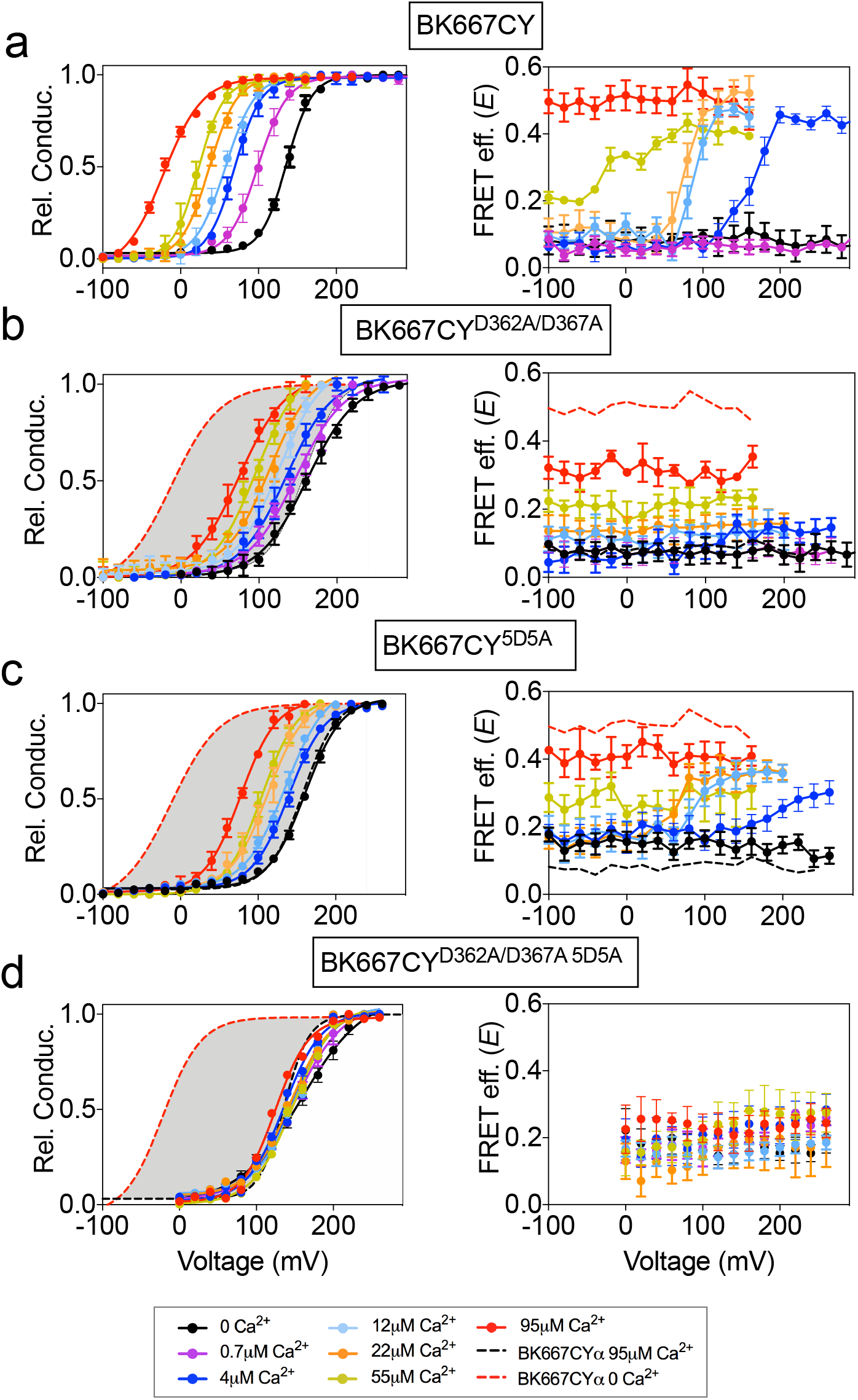
Voltage dependence of gating ring rearrangements is associated to activation of the RCK1 binding site. G-V (left panels) and *E*-V curves (right panels) obtained simultaneously at several Ca^2+^concentrations from **(a)** the BK667CY construct, **(b)** mutation of the RCK1 high-affinity site (D362A/D367A), **(c)** mutation of the Ca^2+^ bowl (5D5A), or **(d)** both (D362A/D367A 5D5A). Note that the voltage dependence of the *E* signal is only abolished after mutating the RCK1 high-affinity binding site (b) or both (d). Data corresponding to each Ca^2+^ concentration are color-coded as indicated in the legend at the bottom. Solid curves in the G-V graphs represent Boltzmann fits. For reference, grey shadows in **a-d** left panels represent the full range of G-V curves corresponding to non-mutated BK667CY channels from 0 μM Ca^2+^ to 95 μM Ca^2+^ (indicated with colored dashed lines). Data points and error bars represent average ± SEM (*n*=3-14). Data in **a** and **b** are taken from (31) and (30), respectively.

As described previously, the G-V curves show a significantly reduced shift to more negative potentials when Ca^2+^ is increased, as compared to the non-mutated BK667CY (**Fig. 1a-b**, left panels). Specific activation of the Ca^2+^ bowl renders a smaller change in *E* values, which are not voltage-dependent within the voltage range tested (**Fig. 1b**, right panel). To test the effect of eliminating the RCK2 Ca^2+^ binding site -the Ca^2+^ bowl- we mutated five aspartates to alanines (5D5A) (28). As expected, activation of only the RCK1 domain by Ca^2+^ reduced the Ca^2+^-dependent shift in the GV curves (**Fig. 1c**, left panel). Even though the extent to which the *E* values changed with Ca^2+^ was reduced (**Fig. 1c**), there was a persistent voltage dependence equivalent to that detected in the non-mutated channel (**Fig. 1c**, right panel) (31). This effect seems not to be attributable to Ca^2+^ binding to unknown binding sites in the channel, since the double mutation of the RCK1 and RCK2 sites abolishes the change in the FRET signal (**Fig. 1d**). Altogether, these results indicate that the voltage-dependent component of the gating ring conformational changes triggered by Ca^2+^ in the BK667CY construct depends on activation of the RCK1 binding site. Because the gating ring is not within the transmembrane region, it is not expected to be directly influenced by the transmembrane voltage. Therefore, the voltage-dependent FRET signals must be coupled to the dynamics of the gate region associated with the opening and closing of the channel and/or those of the voltage sensor domain.

### The voltage-dependent conformational changes of the gating ring are not related to the opening and closing of the pore domain

To test whether the voltage-dependent FRET signals relate to the opening and closing of the channel (intrinsic gating) we used two modifications of BK channel function in which the relative probability of opening is shifted in the voltage axis, yet the actual dynamics of the intrinsic gating are expected to be unaltered (**Fig. 2b**). We reasoned that if the voltage-dependent FRET signals of the gating ring are coupled to the opening and closing, they should follow a similar displacement with voltage. The first BK channel construct is the α subunit including the single point mutation F315A, which has been described to shift the voltage dependence of the relative conductance of the channel to more positive potentials, by uncoupling the voltage sensor activation from the gate opening (**Fig. 2c**) (35). **Fig. 2d** shows the relative conductance and *E vs*. voltage for the BK667CY^F315A^ mutant at various Ca^2+^concentrations. Our results show that the shift of the relative probability of opening to more positive potentials (**Fig. 2d**, left panel) does not lead to changes in the voltage dependence of the gating ring FRET signals (**Fig. 2d**, right panel).

**Fig. 2.**
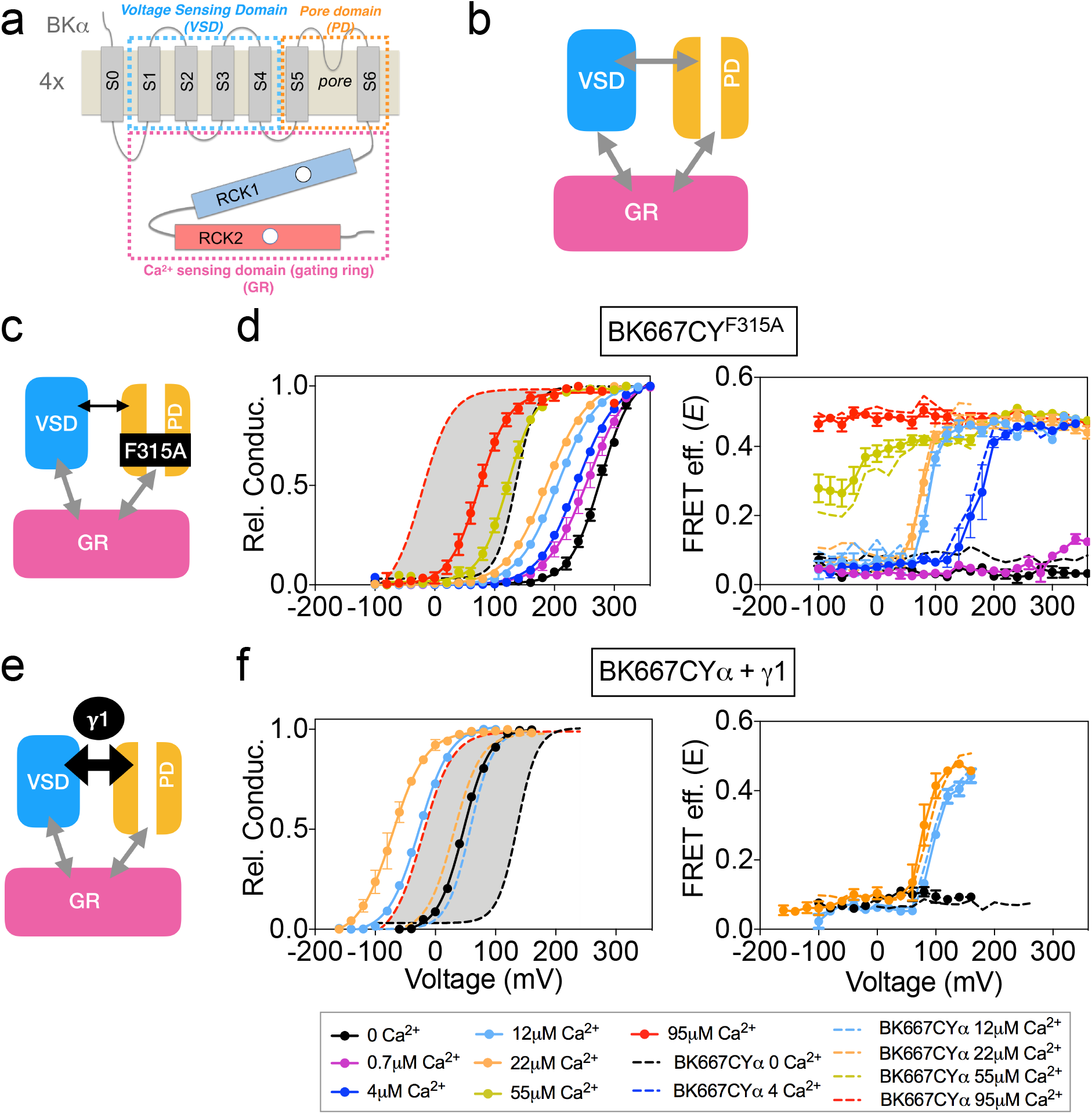
Modification of the voltage dependence of gate opening does not affect the gating ring voltage-dependent conformational changes. **(a)** Topology of the BKα subunit where the voltage sensing domain (VSD), Ca^2+^ sensing domain (gating ring, GR) and pore domain (PD) are indicated by colored dashed lines boxes (see main text for a full description). **(b)** The three BK functional modules (VSD, PD, GR), schematically represented as colored boxes, interact allosterically. **(c)** Diagram representing the main effect of the F315A mutation, which is the uncoupling of the VSD to the PD. **(d)** G-V (left panel) and *E*-V curves (right panel) obtained simultaneously at several Ca^2+^concentrations after mutation of the F315 site to alanine (BK667CY^F315A^). **(e)** The interaction with the γ1 subunit favors the VSD-PD coupling mechanism **(f)** G-V (left) and *E*-V curves (right) of BK667CY α subunits co-expressed with γ1 subunits. In all panels, data corresponding to each Ca^2+^ concentration are color-coded as indicated in the bottom legend. Colored dashed lines represent the G-V and *E*-V curves corresponding to BK667CYα channels (30, 31). The solid curves in the G-V graphs represent Boltzmann fits. The full range of G-V curves from 0 μM Ca^2+^ to 95 μM Ca^2+^ from BK667CY is represented as a grey shadow in left panels **d** and **f**, for reference. Data points and error bars represent average ± SEM (*n*=3-6).

The second modification of BK function consisted in co-expressing the wild type α subunit with the auxiliary subunit γ1 (36–39). In this case, the relative probability of opening is shifted to more negative potentials by increasing the coupling between the voltage sensor and the gate of the channel (**Fig. 2e**). This construct adds the advantage of representing a physiologically relevant modification of channel gating. **Fig. 2f** shows the relative conductance and *E vs*. voltage in oocytes co-expressing the BK667CYα and γ1 at voltages ranging from −160 to +260 mV, with three [Ca^2+^] concentrations: nominal 0, 12 μM and 22 μM. As expected, the presence of the γ1 subunit drives the relative conductance curves to more negative potentials (**Fig. 2f**, left panel) compared to the values obtained without γ1 (**Fig. 2f**, dashed lines). Remarkably, the change in the voltage dependence of the relative conductance induced by γ1 does not alter the simultaneously recorded FRET signals (**Fig 2f**, right panel), which remains indistinguishable from that recorded with BK667CYα (**Fig. 2f**, dashed lines).

### The dynamics of the VSD are directly reflected in the gating ring conformation

In the allosteric model of BK channel function, Ca^2+^ binding to the Ca^2+^ bowl is weakly coupled to the voltage sensor activation (reflected in the low value of the allosteric constant E) (3). Nevertheless, the model does not discount some level of interaction between the VSD and the gating ring, so we decided to explore if the voltage dependence of the gating ring conformational change is attributable to the voltage sensor activation. To this end, we modified the voltage dependence of the voltage sensor activation by co-expression with β auxiliary subunits or by introducing specific mutations in the VSD (**Fig. 3** and **Fig. 4**). The effects of co-expressing BK α subunit with the four different types of auxiliary β subunits have been extensively studied (37, 40–49). β1 subunit has been previously proposed to alter the voltage sensor-related voltage dependence, as well as the intrinsic opening of the gate and Ca^2+^ sensitivity (**Fig. 3a**) (42–44, 46, 47, 50). Recordings from BK667CYα co-expressed with β1 subunits reveal the expected modifications in the voltage dependence of the relative conductance, i.e. an increase in the apparent Ca^2+^ sensitivity (**Fig. 3b**, left panel) (42–44, 46, 47, 51). In addition it has been reported that β1 subunit alters the function of the VSD (42, 50). Notably, the *E*-V curves are shifted to more negative potentials (**Fig. 3b**, right panel), similarly to the described modification (50). The structural determinants of the β1 subunit influence on the VSD reside within its N-terminus, which has been shown by engineering a chimera between the β3b subunit (which does not influence the VSD) and the N-terminus of the β1 (β3bNβ1) (50). We recapitulated this strategy. First, we co-expressed BK667CY α subunits with β3b and observed the expected inactivation of the ionic currents at positive potentials (**Supplementary Figure 1**) (52–54). The relative open probability of this complex is like BK667CYα alone, except that at extreme positive potentials the values of relative conductance decrease due to inactivation (**Supplementary Figure 1b**, left panel) (45, 54). The values of *E* vs V remained comparable to those observed for BK667CYα (**Supplementary Figure 1**, right panel). We then co-expressed the β3bNβ1 chimera (50) with BK667CYα (**Fig. 3c**). This complex did not modify the relative conductance *vs*. voltage relationship (**Fig. 3d**, left panel) as compared with BK667CYα alone (**Fig. 3d**, grey shadow). On the other hand, while the magnitude of the FRET change is the same as in BK667CYα, the voltage dependence of *E* values at [Ca^2+^] of 4 μM, 12 μM and 22 μM shifted to more negative potentials compared to the values of BK667CYα alone (**Fig. 3d**, right panel, compare dashed to solid lines). Altogether, these results indicate that the alteration of the voltage dependence of the voltage sensor induced by the amino terminal of β1within the β3bNβ1 chimera underlies the modification of the voltage dependence of the gating ring conformational changes, reinforcing the hypothesis that this voltage dependence is directly related to VSD function.

**Fig. 3.**
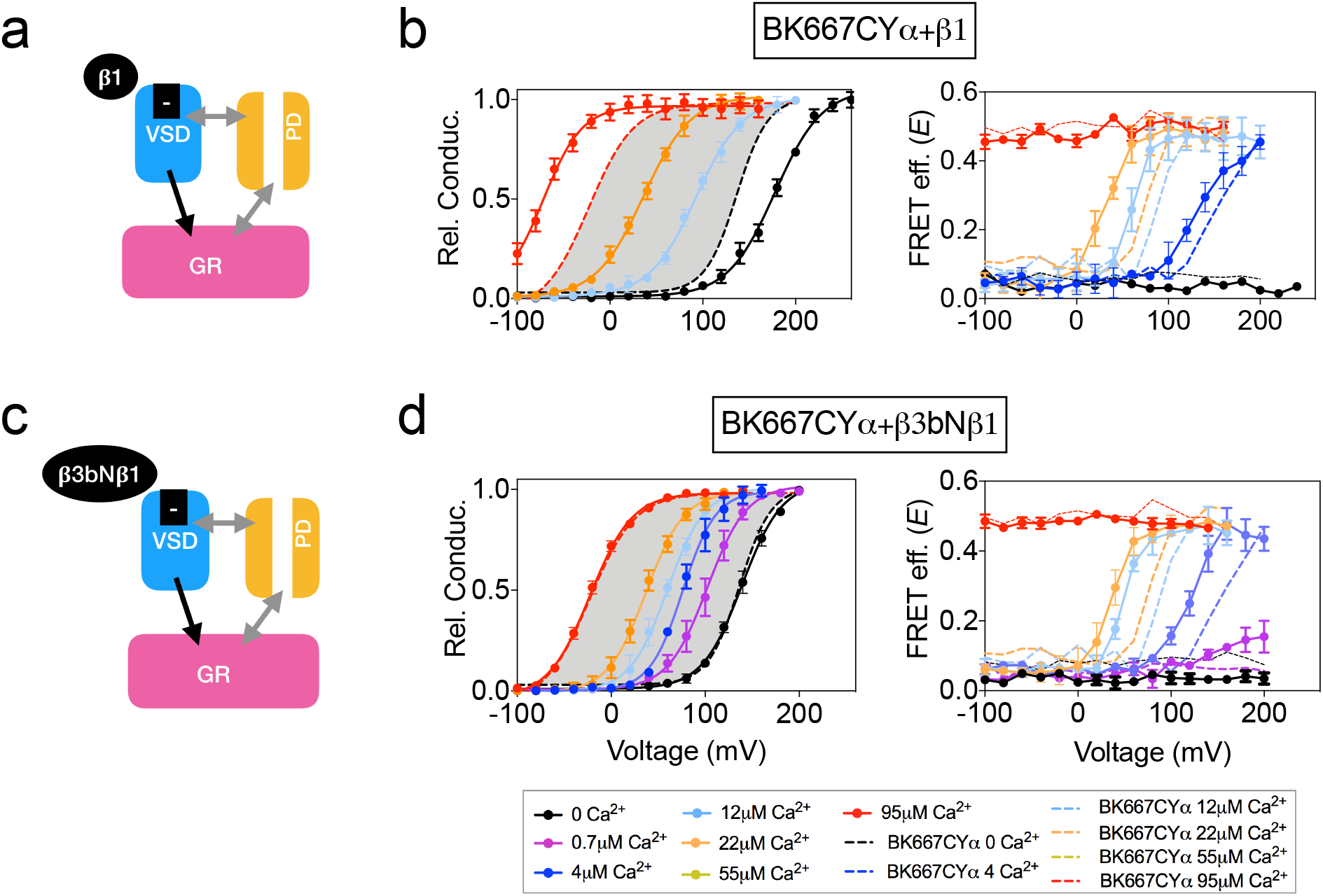
Co-expression with β subunits. **(a)** β1 subunits have been shown to directly regulate VSD function, shifting V_h(j)_ to more negative values **(b)** Left panel, G-V curves obtained at several Ca^2+^concentrations after co-expression of BK667CY with the β1 subunit, which induces a leftward shift in the *E*-V curves obtained simultaneously (right). **(c)** β3bNβ1 chimeras produce similar effects to β1 on VSD function, since they retain the N-terminal region of β1 (50). **(d)** G-V (left) and *E*-V curves (right) of BK667CY α subunits co-expressed with the β3bNβ1 chimera. Data corresponding to each Ca^2+^ concentration are color-coded as indicated in the legend at the bottom. Colored dashed lines represent the G-V and *E*-V curves corresponding to BK667CYα channels (30, 31). The solid curves in the G-V graphs represent Boltzmann fits. The full range of G-V curves from 0 μM Ca^2+^ to 95 μM Ca^2+^ from BK667CY is represented as a grey shadow in left panels **b** and **d**, for reference. Data points and error bars represent average ± SEM (*n*=3-10).

**Fig. 4.**
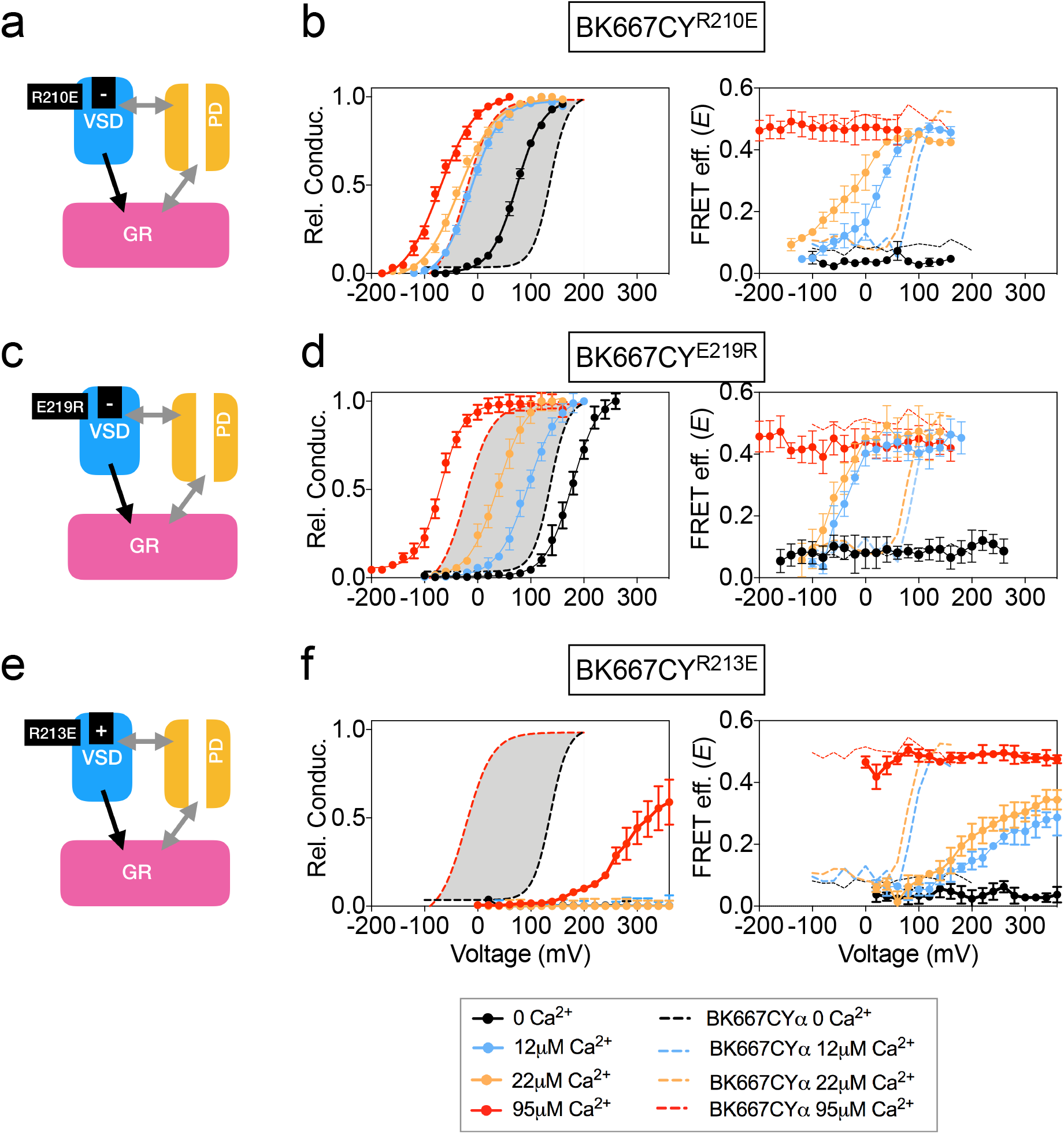
Mutation of charged residues of BK VSD. VSD activation was altered by mutation of charged residues in the VSD that modify its voltage of half activation, V_h(j)_ **(a)** The R210E mutation induces a negative shift of V_h(j)_ **(b)** G-V (left panel) and *E*-V curves (right panel) obtained simultaneously from constructs BK667CY containing the R210E mutation at several Ca^2+^concentrations. **(c)** The E219R mutation produces a negative shift of V_h(j)_ **(d)** G-V (left panel) and *E*-V curves (right panel) obtained simultaneously from constructs BK667CY containing the E219R mutation at several Ca^2+^concentrations. **(e)** The R213E mutation induces a large positive shift of V_h(j)_ values. **(f)** G-V (left panel) and *E*-V curves (right panel) obtained simultaneously from constructs BK667CY containing the R213E mutation at several Ca^2+^concentrations. Data corresponding to each Ca^2+^ concentration are color-coded as indicated in the bottom legend. Colored dashed lines represent the G-V and *E*-V curves corresponding to non-mutated BK667CYα channels (30, 31). The solid curves in the G-V graphs represent Boltzmann fits. The full range of G-V curves from 0 μM Ca^2+^ to 95 μM Ca^2+^ from BK667CY is represented as a grey shadow in left panels **b, d** and **f**, for reference. Data points and error bars represent average ± SEM (*n*=3-10).

VSD activation can also be altered by introducing single point mutations that modify the voltage of half activation of the voltage sensor, V_h_(j). This parameter is determined by fitting data to the HA allosteric model (19) or directly from gating current measurements (55). Mutations of charged amino acids on the VSD have been reported to produce different modifications in the V_h_(j) values. In some cases, other parameters related to BK channel activation are additionally affected by the mutations. Mutation R210E shifts the V_h_(j) value from +173 mV to +25 mV at 0 Ca^2+^ in BK channels (**Fig. 4a**) (19). Consistently, introduction of this mutation in BK667CYα (BK667CY^R210E^) caused a shift of the relative conductance *vs*. voltage dependence towards more negative potentials (**Fig. 4b**, left panel) as compared to BK667CY (**Fig. 4b**, left panel, grey shadow). Simultaneously measured *E* values showed a negative shift in the voltage dependence of the FRET signal at intermediate Ca^2+^ concentrations (**Fig. 4b**, right panel). Mutation E219R had been previously shown to produce a large negative shift in V_h_(j) from +150 mV to +40 mV (ΔV_h_(j) = −110 mV; **Fig. 4c**), additionally modifying the Ca^2+^ sensitivity and the coupling between the VSD and channel gate (55). As previously reported, BK667CY^E219R^ showed modified relative conductance *vs*. voltage relationships at different Ca^2+^ concentrations (**Fig. 4d**, left panel) (55). In addition, this construct revealed shift to more negative potentials in the *E vs*. voltage dependence at intermediate Ca^2+^ concentrations (12 μM and 22 μM Ca^2+^; **Fig 4d**, right panel), paralleling the reported negative shift in V_h_(j) (19, 55). Since mutations displacing the V_h_(j) to more negative potentials induce equivalent shifts in the voltage dependence of the gating ring motion (measured as *E*), we tested if other mutations previously reported to induce positive shifts on V_h_(j) (19) were also associated to changes of the *E*-V curves in the same direction. As shown by Ma et al., the largest effect on V_h_(j) is induced by the R213E mutation, producing a shift of ΔV_h_(j)=+337mV (**Fig. 4e**) (19). The BK667CY^R213E^ construct showed a significant shift in the voltage dependence of the relative conductance to more positive potentials (**Fig. 4f**, left panel). Notably, this effect was paralleled by a large displacement in the *E vs*. voltage dependence towards more positive potentials (**Fig. 4f**, right panel). Taken together, our data show that modifications of the V_h_(j) values caused by mutating the VSD charged residues are reflected in equivalent changes in the voltage dependence of the gating ring conformational rearrangements, which occur in analogous directions and with proportional magnitudes at intermediate Ca^2+^ concentrations.

All these results on the VSD modifications and their corresponding changes in FRET signals support the existence of a direct coupling mechanism between the VSD function and the gating ring conformational changes.

### Parallel alterations of the voltage dependence of VSD function and gating ring motions by selective activation of the RCK1 binding site

We have previously shown that specific interaction of Cd^2+^ with the RCK1 binding site leads to activation of the BK channel, which is accompanied by voltage-dependent changes in the *E* values at intermediate Cd^2+^ concentrations of 10 μM and 30 μM (30). To further assess the role of the RCK1 binding site activation in the voltage dependence of the gating ring motions, we studied activation by Cd^2+^ of selected BK667CY VSD mutants (**Fig. 5**). Addition of Cd^2+^ to the BK667CY^E219R^ mutant (**Fig. 5a**) shifted the voltage dependence of *E* towards more negative potentials at intermediate Cd^2+^ concentrations (10 μM and 30 μM; **Fig. 5b**) when compared to non-mutated BK667CY (**Fig. 5b**; dashed lines). This change in the *E*-V curves induced by selective activation of the RCK1 binding site with Cd^2+^ paralleled the large negative shift (ΔV_h_(j) = −110 mV) previously reported with the E219R mutant BK channels (19, 55). We also tested Cd^2+^ activation in the mutant BK667CY^R201Q^, which shifts the V_h_(j) parameter by 47 mV towards positive potentials (**Fig. 5c**) (19). Addition of Cd^2+^ rendered right-shifted *E* vs. voltage relationships (**Fig.5d**, right panel), following the direction of the predicted V_h_(j) shift described for this mutant BK channel (19). Finally, addition of Cd^2+^ to the BK667CY^F315A^ construct (**Fig. 5e**) (35) did not have any effect on the *E*-V relationship (**Fig. 5f**). These results are consistent with a mechanism in which specific binding of Cd^2+^ to the RCK1 binding site allows voltage-dependent conformational changes in the gating ring that are directly related to VSD activation.

**Fig. 5.**
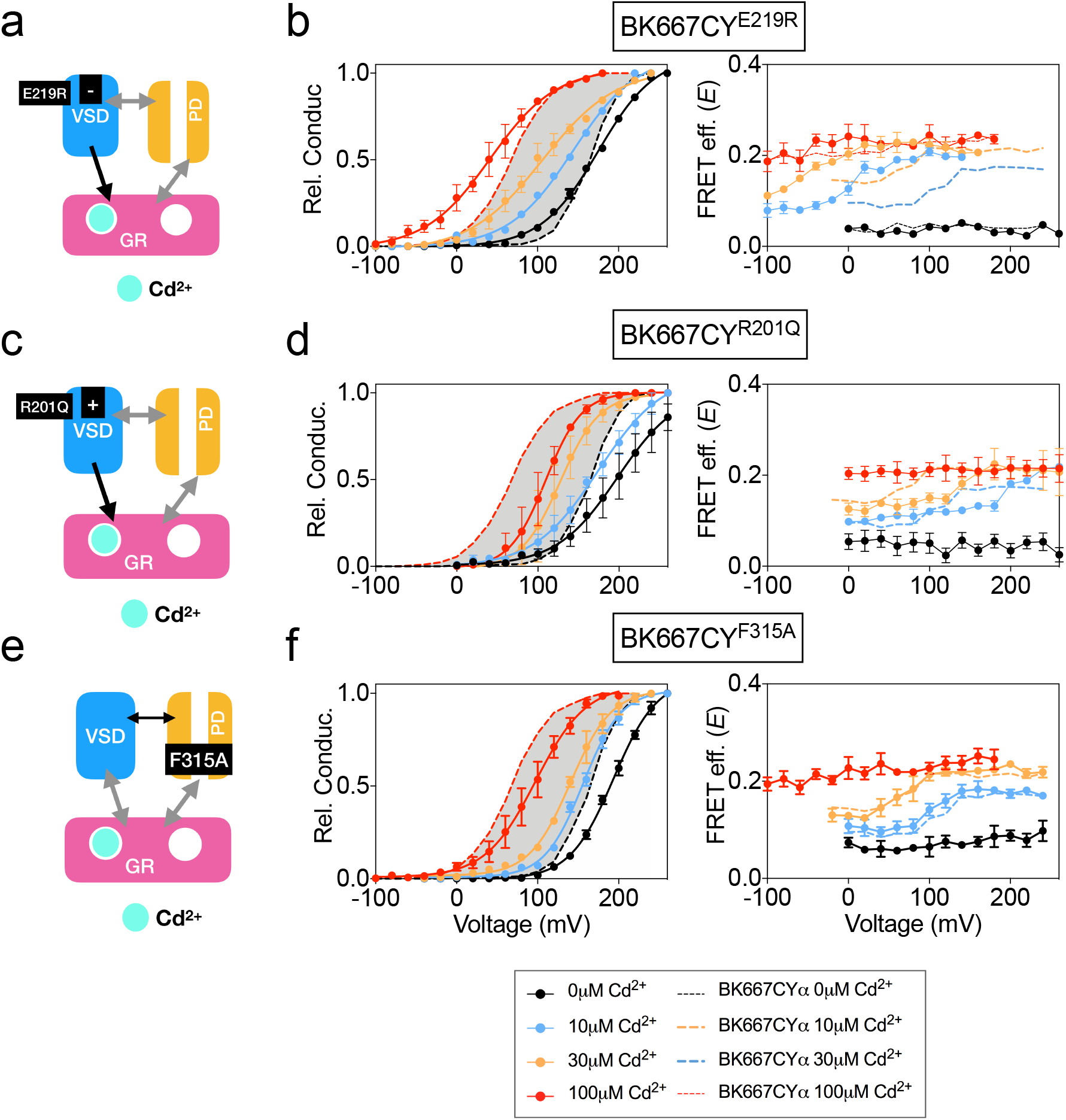
Voltage dependence of gating ring rearrangements after specific activation of RCK1 high-affinity binding site by Cd^2+^. **(a)** Effect of the VSD E219R mutation on the selective activation of RCK1 by Cd^2+^. **(b)** G-V (left panels) and *E*-V curves (right panels) obtained simultaneously at several Ca^2+^concentrations from constructs BK667CY^E219R^. **(c)** VSD R201Q mutation induces a positive shift of V_h(j)_ **(d)** G-V (left panels) and *E*-V curves (right panels) obtained simultaneously at several Ca^2+^concentrations from constructs BK667CY^R201Q^ **(e)** Effect of the F315A mutation on the selective activation of RCK1 by Cd^2+^. **(f)** G-V (left panels) and *E*-V curves (right panels) obtained simultaneously at several Ca^2+^concentrations from constructs BK667CY^F315A^. Data corresponding to each Ca^2+^ concentration are color-coded as indicated in the legend at the bottom. Colored dashed lines represent the G-V and *E*-V curves corresponding to BK667CYα channels (30, 31). The solid curves in the G-V graphs represent Boltzmann fits. The full range of G-V curves from 0 μM Cd^2+^ to 100 μM Cd^2+^ corresponding to non-mutated BK667CY is represented as a grey shadow in left panels **b, d**, and **f**, for reference. Data points and error bars represent average ± SEM (*n*=3-10).

### Voltage dependence of Ba^2+^-induced gating ring movement is related to function of the channel gate

Ca^2+^, Mg^2+^ and Ba^2+^ bind to the Ca^2+^ bowl and trigger conformational changes of the gating ring region (30). However, the effects of these ions on BK function and gating ring motions are fundamentally different. Notably, Ba^2+^ induces a rapid blockade of the BK current after a transient activation that is measurable at low Ba^2+^ concentrations (29, 30) (**Fig. 6a**). In addition, we previously showed that the gating ring conformational motions induced by Ba^2+^ show a voltage-dependent component, which is not observed when Ca^2+^ or Mg^2+^ bind to the Ca^2+^ bowl (30, 31) (**Fig. 6b**). We combined mutagenesis with the cation-specific activation strategy to identify the structural source of the voltage dependence in Ba^2+^-triggered gating ring motions. In this case, alteration of VSD function by mutating charged residues (**Fig. 6c** and **6e**) was not reflected in any change of the *E vs*. voltage relationships, as shown in **Fig. 6d** and **6f** for constructs BK667CY^R210E^ and BK667CY^R213E^, respectively. These results indicate that the voltage dependence of Ba^2+^-induced gating ring conformational changes, unlike those induced by Ca^2+^ and Cd^2+^ through activation of the RCK1 binding site, may not be related to VSD activation. To further test this hypothesis, we studied the effect of Ba^2+^ on BK667CY channels containing the F315A mutation (**Fig. 6g**) (35). As shown in **Fig. 6h**, the *E* values reached similar levels to those of non-mutated BK667CY channels at saturating Ba^2+^ concentrations. However, at intermediate concentrations of Ba^2+^ the *E*-V curves were shifted towards more positive potentials when compared with BK667CY channels (**Fig. 6h**, dashed line). These results suggest that the voltage-dependent component of the conformational changes triggered by Ba^2+^ binding to the Ca^2+^ bowl are not directly related to VSD activation, but rather to the function of the channel gate.

**Fig. 6.**
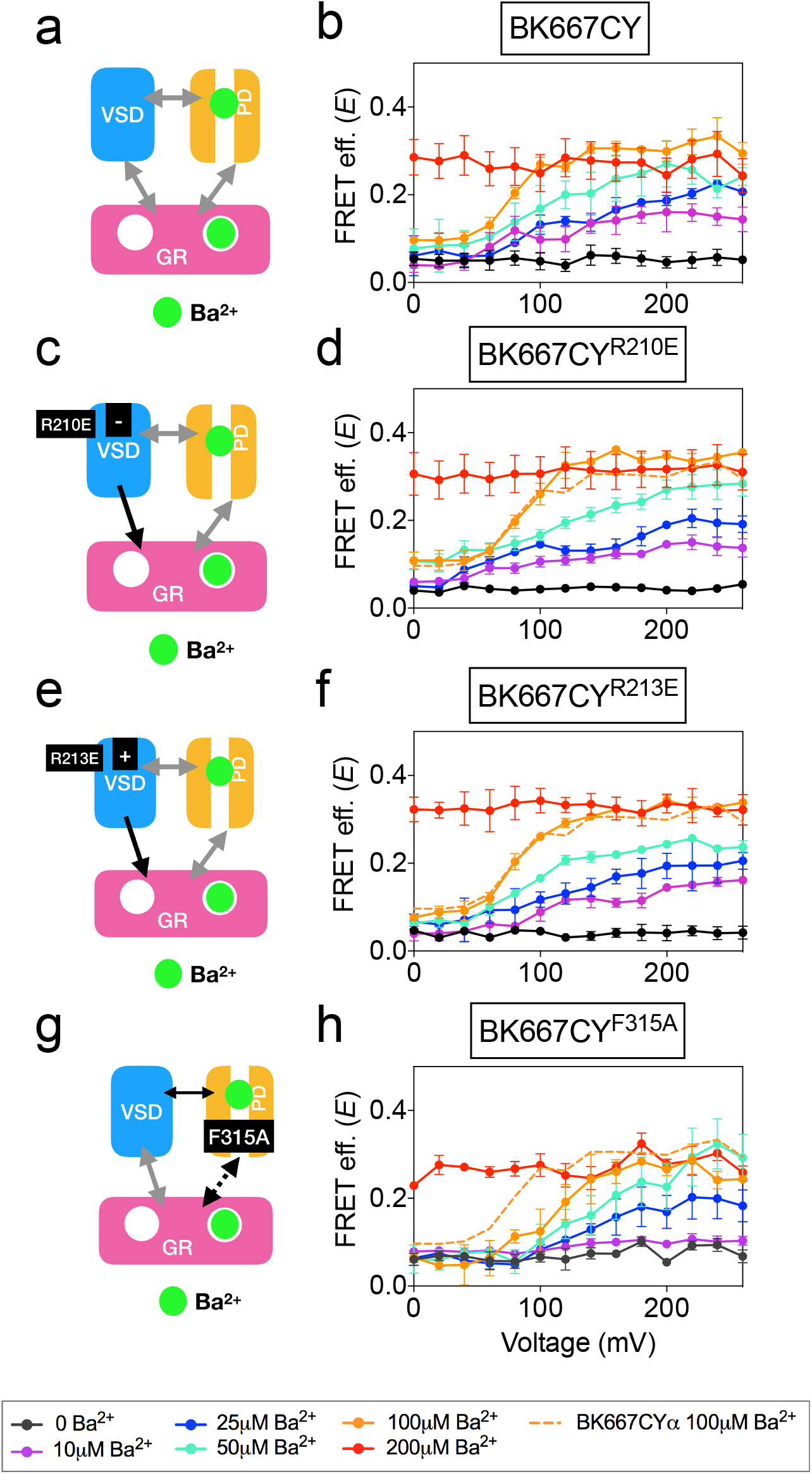
Voltage dependence of gating ring movements triggered by Ba^2+^. **(a)** The RCK2 site is selectively activated by Ba^2+^, which additionally induces pore block. **(b)** FRET efficiency (*E*) data obtained at several Ba^2+^ concentrations from BK667CY constructs (30). **(c)** Effect of the VSD R210E mutation after selective activation of the RCK2 binding site by Ba^2+^. **(d)** *E*-V curves obtained at several Ba^2+^ concentrations from BK667CY^R210E^ constructs. **(e)** Effect of the VSD R213E mutation after selective activation of the RCK2 binding site by Ba^2+^ **(f)** *E*-V curves obtained at several Ba^2+^ concentrations from BK667CY^R213E^ constructs. **(g)** Effect of the F315A mutation after selective activation of the RCK2 binding site by Ba^2+^ **(h)** *E*-V curves obtained at several Ba^2+^ concentrations from BK667CY^F315A^ constructs. Data corresponding to each Ba^2+^ concentration are color-coded according to the legend at the bottom. For reference, the curve corresponding to 100 μM Ba^2+^ from the BK667CY construct shown in **(b)** is also shown as a colored dashed line in panels **b, d, f** and **h**. Data points and error bars represent average ± SEM (*n*=3-7).

## DISCUSSION

Using fluorescently labeled BKα subunit constructs reporting protein dynamics between the RCK1 and RCK2 domains, we previously demonstrated that the channel high-affinity binding sites can be independently activated by different divalent ions, inducing additive rearrangements of the gating ring measured as changes in the FRET efficiency values, *E* (30, 31). Further, the effects of Ca^2+^, Cd^2+^ and Ba^2+^ on the *E* values showed a voltage-dependent component, for which we could not provide a structural basis. Voltage dependence of Ca^2+^-induced rearrangements seemed to be specifically related to RCK1 activation, since only the mutation of that binding site resulted in voltage-independent *E* signals ((30) and **Fig. 1**). One possibility to explain this result is the existence of direct structural interactions of the RCK1 domain and the VSD. Interestingly, recently obtained cryo-EM full BK structure from *Aplysia californica* revealed the existence of specific protein-protein interfaces formed by the amino terminal lobes of the RCK1 domains facing the transmembrane domain and the VSD/S4-S5 linkers (22). According to the structural data, gating of the channel by Ca^2+^ was proposed to be mediated, at least partly, by displacement of these interfaces causing the VSD/S4-S5 linkers to move, contributing to pore opening (15, 22). Our work now provides functional data supporting this mechanism. Our data show that mutations altering the voltage dependence of BK VSD are reflected in the voltage dependence of the gating ring movements triggered by activation of the RCK1 binding site by Ca^2+^ or Cd^2+^. Mutations altering VSD function by inducing large leftward shifts in the V_h_(j) values (19, 55) strongly correlate with negative shifts in the voltage dependence of the *E* signals. Likewise, mutations inducing positive shifts in the VSD voltage dependence of the voltage sensor function are reflected in *E*-V shifts towards more positive membrane voltages. The strong correlation between the voltage of half maximum activation of the voltage sensor V_h_(j) and that of the *E*-V curves suggests the existence of an interaction between the VSD and the gating ring. These results are further supported by the effect of auxiliary subunits. We have tested the effect of γ1, β1 and β3b on the voltage dependence of the *E* signals. Within these three subunits, only β1 has been proposed to alter the voltage dependence of VSD function. Consistently with our hypothesis, γ1 and β3b induce the expected effects on the G-V curves of BK667CY channels, without altering the *E*-V curves, whereas β1 induces a leftward shift in the *E*-V curves. This shift on the voltage dependence of the *E* signal is preserved when BK667CY channels are coexpressed with the β3bNβ1 chimera, which affects VSD function without altering other parameters related to gating (50). Therefore, the voltage-dependent component of the *E* signal seems to be related strictly to VSD function, as further demonstrated by the lack of effect of the γ1 subunit, which has been shown to shift the voltage dependence of gate opening by enhancing the allosteric coupling of voltage sensor activation without affecting VSD operation (36). Similar lack of effect on the voltage dependence of *E* is observed after introducing into BK667CY channels the mutation F315A, which has been also associated to uncoupling VSD function from pore opening (35).

A puzzling result from our previous study was the observation that Ba^2+^ binding to the Ca^2+^ bowl triggers voltage-dependent conformational changes (30). Even though we still do not know the mechanisms of this unique response to Ba^2+^, here we learned that it is not related to the dynamics of VSD, but rather influenced by perturbations affecting the opening and closing of the channel at the pore domain. A possible explanation for this result is that Ba^2+^ block of the permeation pathway (16, 29, 56) is somehow transmitted allosterically to the gating ring. Alternatively, there could be a direct allosteric interaction between the intrinsic gating region and the divalent binding site in RCK2.

Irrespectively of the fluorescent construct (31) or the divalent ion used to activate the BK channel (30), we have consistently observed that the conformational changes monitored as changes in the FRET efficiency are not strictly coupled to the intrinsic gating of the channel. In this study, we have found that the consequences of the voltage dependence of the intrinsic gating by manipulations of the VSD and the pore region are paralleled by the FRET efficiencies. These results rule out the possibilities that FRET signals derive from conformational changes in an unknown Ca^2+^ binding site or that they are completely uncoupled to the intrinsic gating.

In conclusion, our functional data show a strong correlation between the VSD function and the RCK1 conformational changes, suggesting a transduction mechanism from ion binding to change the channel activation. This transduction mechanism is in agreement with the existence of structural interactions between the RCK1 domain and the VSD. The strong correlation between VSD function and the RCK1 conformational changes is not observed between RCK2 and VSD, suggesting the existence of a different transduction mechanism that may include an indirect mechanism through the RCK1 or RCK1-S6 linker.

## Methods

### Molecular biology and heterologous expression of tagged channels

Fluorescent BK α subunits were labelled with CFP or YFP using a transposon based insertion method (57). Subunits labelled in the position 667 were subcloned into the pGEMHE oocyte expression vector (58). RNA was transcribed *in vitro* with T7 polymerase (Ambion, Thermo Fisher Scientific, Waltham, USA), and injected at a ratio 3:1 of CFP: YFP into *Xenopus laevis* oocytes, giving a population enriched in 3CFP:1YFP labelled tetramers (BK667CY) (30, 31). Individualized Oocytes were obtained from *Xenopus laevis* extracted ovaries (Nasco, Fort Anderson, WI, USA). Neutralization of the Ca^2+^ bowl was achieved by mutating five consecutive aspartate residues to alanines (5D5A: 894-899) (28) on the BK667CY background. Elimination of RCK1 high-affinity Ca^2+^ sensitivity was achieved by double mutation D362A and D367A (2, 4, 27). Elimination of the Mg^2+^ binding site was archived with mutation D99A (26). Mutations were performed using standard procedures (Quickchange, Agilent Technologies, Santa Clara, USA).

### Patch-clamp fluorometry and FRET

Borosilicate pipettes with a large tip (0.7-1 MΩ in symmetrical K^+^) were used to obtain inside-out patches excised from *Xenopus laevis* oocytes expressing BK667CY. Currents were recorded with the Axopatch 200B amplifier and Clampex software (Axon Instruments, Molecular Devices, Sunnyvale, USA). Recording solutions contained (in mM): pipette, 40 KMeSO_3_, 100 N-methylglucamine-MeSO_3_, 20 HEPES, 2 KCl, 2 MgCl_2_, 100 μM CaCl_2_ (pH 7.4); bath solution, 40 KMeSO_3_, 100 N-methylglucamine-MeSO_3_, 20 HEPES, 2 KCl, 1 EGTA, and MgCl_2_ or BaCl_2_ to give the appropriate divalent concentration previously estimated using Maxchelator software (maxchelator.standford.edu) (59). Solutions containing Cd^2+^ were prepared with a bath solution containing KF instead of K-Mes to precipitate the contaminant Ca^2+^ previously to the administration of the proper concentration of CdCl_2_ estimated with Maxchelator. Solutions containing different ion concentrations were exchanged using a fast solution-exchange system (BioLogic, Claix, France).

Simultaneous fluorescent and electrophysiological recordings were obtained as previously described (30, 31). Conductance-voltage (G-V) curves were obtained from tail currents using standard procedures. Conformational changes of the gating ring were tracked as intersubunit changes of the FRET efficiency between CFP and YFP as previously reported (30, 31). Analysis of the FRET signal was performed using emission spectra ratios. We calculated the FRET efficiency as *E*=(RatioA-RatioA_0_)/(RatioA_1_-RatioA_0_), where RatioA and RatioA_0_ are the emission spectra ratios for the FRET signal and the control only in the presence of acceptor respectively (60); RatioA_1_ is the maximum emission ratio that we can measure in our system (30, 31). This value of *E* is proportional to FRET efficiency (60). The *E* value showed is an average of the *E* value corresponding to each tetramer present in the membrane patch and represent an estimation of the distance between the fluorophores located in the same position of the four subunits of the tetramer.

## Acknowledgments

MH and PM were supported by the intramural section of the National Institutes of Health (NINDS). TG was funded by the Spanish Ministry of Economy and Competitivity (grants SAF2013-50085-EXP and RyC-2012-11349) and the European Research Council (ERC) under the European Union’s Horizon 2020 research and innovation programme (grant agreement 648936). We thank Deepa Srikumar for technical assistance. The γ1 clone and the β1β3b chimera were kind gifts from Chris Lingle and Ramon Latorre, respectively.

---

**Supplementary Figure 1.**
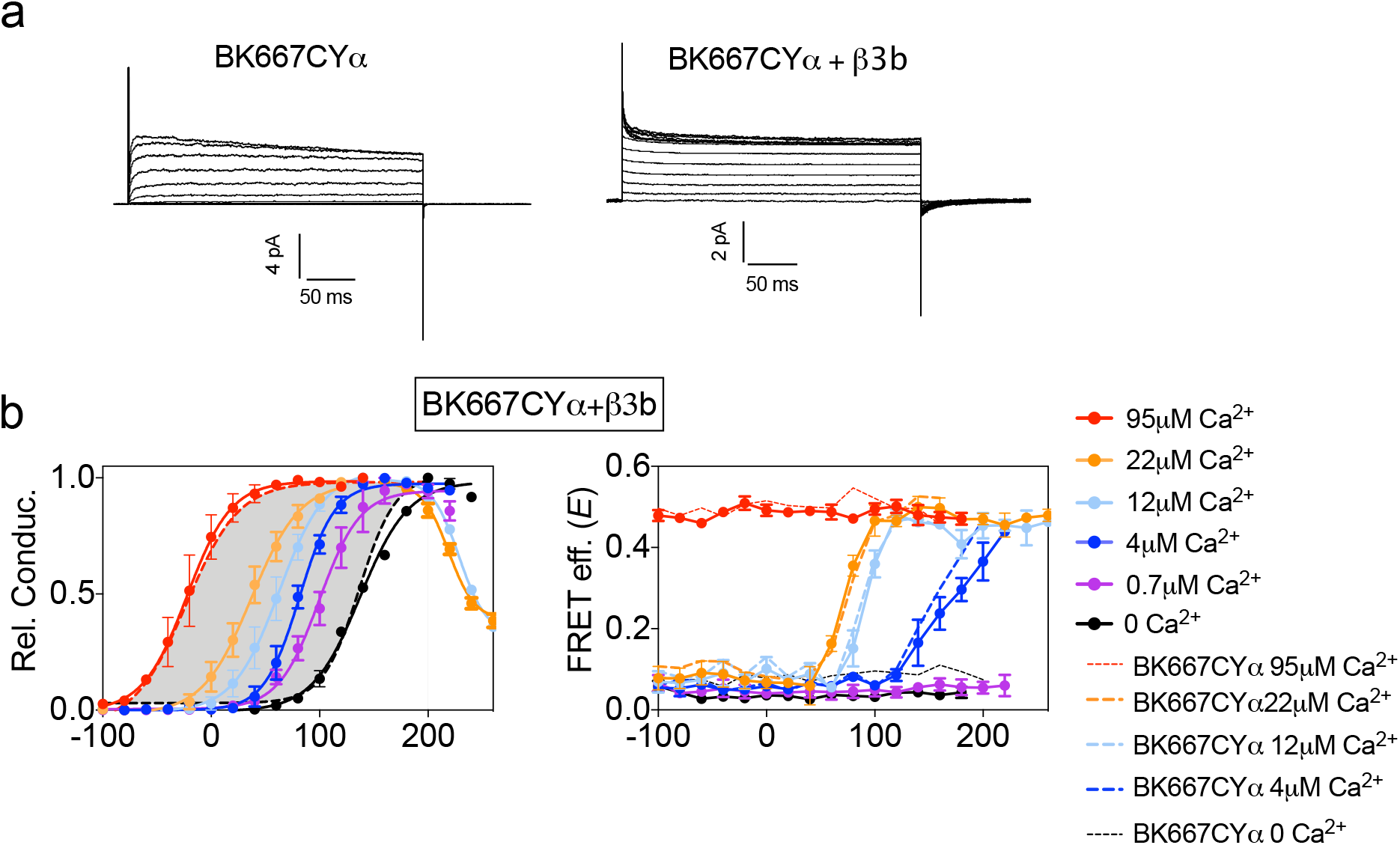
Co-expression with β *subunits*. **(a)** Representative currents obtained after applying depolarizing pulses to inside-out patches expressing BK667CYα (left) or BK667CYα + β3b channels, in the presence of 12 μM Ca^2+^. **(b)** Left panel, G-V curves obtained at several Ca^2+^concentrations after co-expression of BK667CY with β3b subunits, inducing no appreciable changes in the *E*-V curves obtained simultaneously (right).

